# A link between mitochondrial gene expression and life stage morphologies in *Trypanosoma cruzi*

**DOI:** 10.1101/798421

**Authors:** Roger Ramirez-Barrios, Emily K. Susa, Sean P. Faacks, Charles K. Liggett, Sara L. Zimmer

**Affiliations:** University of Minnesota Medical School, Duluth campus, Duluth, MN, USA; University of Minnesota Duluth, Duluth, MN, USA

**Keywords:** Trypanosomiasis, Organelles, Gene Expression, RNA editing, Life Cycle Stages, Ribonuclease

## Abstract

The protozoan *Trypanosoma cruzi* has a complicated dual-host life cycle, and starvation can trigger transition from the replicating insect stage to the mammalian-infectious nonreplicating insect stage (epimastigote to trypomastigote differentiation). Abundance of some mature RNAs derived from its mitochondrial genome increase during culture starvation of *T*. *cruzi* for unknown reasons. Here we examine *T. cruzi* mitochondrial gene expression in the mammalian intracellular replicating life stage (amastigote), and uncover implications of starvation-induced changes in gene expression in insect-stage cells. Mitochondrial RNA levels in general were found to be lowest in actively replicating amastigotes. We discovered that mitochondrial respiration decreases during starvation, despite the previously-observed increases in mitochondrial mRNAs encoding electron transport chain components. Surprisingly, *T. cruzi* epimastigotes in replete medium grow at normal rates when we genetically compromised their ability to perform insertion/deletion editing and thereby generate mature forms of some mitochondrial mRNAs. However, these cells, when starved, were impeded in the epimastigote to trypomastigote transition. Further, they experience a short-flagella phenotype that may also be linked to differentiation. We hypothesize a scenario where levels of mature RNA species or editing in the single *T. cruzi* mitochondrion are linked to differentiation by a yet-unknown signaling mechanism.

## Introduction

In eukaryotes, including many single-celled pathogens, expression of a mitochondrial genome must be coordinated with nuclear gene expression because subunits of mitochondrial electron transport chain (ETC) complexes that participate in oxidative phosphorylation are encoded in both genomes. Proper nuclear-mitochondrial crosstalk is expected to be particularly important for protists as their environments can change rapidly and dramatically, requiring a coordinated response of altered gene expression. However, very little in general is understood about the signals and responses between protist mitochondria and their immediate environment: the cells within which they reside.

Dual-host parasitic trypanosomes are examples of organisms required to rapidly remodel their genomes in response to environmental changes if they are to successfully navigate their life cycle. The best studied trypanosome, *Trypanosoma brucei*, actively respires when replicating in its insect host, but when replicating in the glucose-rich mammalian bloodstream environment is entirely reliant on glycolysis for energy, primarily excreting the pyruvate product of glycolysis rather than extracting additional energy from this metabolic intermediate in its single mitochondrion (Tielens and van Hellemond, 2009). Not surprisingly, mature mRNAs of subunits of ETC complexes III and IV that are derived from the mitochondrial genome are not present at this life stage as they are in *T. brucei*’s insect stage (Feagin and Stuart, 1985; Feagin *et al*., 1988; Bhat *et al*., 1992; Read *et al*., 1994). Many other *T. brucei* mitochondrial gene products are similarly repressed in its replicating mammalian stage, although a few appear to be increased in abundance (Gazestani *et al*., 2017).

While mitochondrial gene expression is well-studied in *T. brucei*, much less is known about how and why it is regulated throughout the life cycle of other trypanosomes, specifically *Trypanosoma cruzi*, the pathogen responsible for American trypanosomiasis. Although *T. cruzi* also has insect (triatomine bug) and mammalian hosts, aspects of its life cycle are very different from that of *T. brucei*. Notably, *T. cruzi* replicates intracellularly rather than extracellularly as *T. brucei* does. Further, *T. cruzi* is known to respire to some extent in all its life stages (Atwood 3rd *et al*., 2005; Goncalves *et al*., 2011; Li *et al*., 2016; Kalem *et al*., 2018). We have begun to close the knowledge gap in understanding *T. cruzi* mitochondrial gene expression by first investigating it in the replicating (epimastigote) and mammalian-infectious (trypomastigote) insect life stages that can be grown and generated in culture (Shaw *et al*., 2016).

We performed our previous study in culture conditions of both replete nutrients and starvation, as starvation is a major signal for differentiation of epimastigotes to trypomastigotes. In an attempt to distinguish between expression responses to starvation cues and differentiation cues, we examined gene expression under abrupt and extreme starvation that does not result in differentiation, and more gradual starvation that results in a subpopulation of cells differentiating to trypomastigotes (~25-50% of the total population). We did not test the protein abundances of trypanosome mitochondrial gene products because they are largely undetectable (Zimmer, 2019).

Trypanosome mitochondrial gene expression studies are confounded by a special nucleotide insertion/deletion type of post-transcriptional RNA editing, specific to this organelle, that is required to generate an open reading frame within many of its 18 coding transcripts. In a previous study, we found that some of the mitochondrial RNAs that do not require editing in *T. cruzi* are increased in abundance upon starvation, a fate that is most likely attributable to an increase in stability. However, an even greater increase in abundance was seen specifically for the edited form of mRNAs that require editing, while pre-edited mRNA levels remained largely constant (Shaw *et al*., 2016). Therefore, some aspect of the editing process is particularly responsive to the starvation signal. Ultimately, the study concluded with two paradoxical findings that we follow up on in the current study. One is that during the slow starvation resulting in culture differentiation, mRNA levels were greatest in starved epimastigotes rather than the trypomastigotes that were part of the same population. The other is that tested nuclear-encoded subunits of ETC complexes were not upregulated during starvation. Both findings call into question what the functional role the increase in mitochondrial mRNAs and rRNAs might be.

Our overall goal is to determine the functional relevance of changing gene expression patterns of the *T. cruzi* mitochondrial genome in the context of its life cycle. Here, we extend our studies to another *T. cruzi* life stage and show that expression of mature mitochondrial RNAs is even lower in the mammalian intracellular replicating stage, the amastigote, than in fed epimastigotes. We then go back and probe whether increased rRNAs and mRNAs in cultured cells undergoing starvation impact ribosome numbers or mitochondrial respiratory output. To explore whether the increases in mature mitochondrial gene expression during starvation are an essential component of life stage differentiation, we genetically depress the ability of *T. cruzi* to perform mitochondrial RNA editing and determine the downstream differentiation-related effects. We end by proposing a model in which during starvation cues *T. cruzi* epimastigotes to stockpile mitochondrial ribosomes and mature mRNAs in preparation for resumption of replication in the amastigote stage that follows that of the metacyclic trypomastigote. We also propose that accumulation of these mRNA species serves as a checkpoint, whereby in the presence of an appropriate abundance of certain mitochondrial mRNAs, the cell is allowed to proceed through differentiation.

## Results

### Mitochondrial gene expression is largely repressed in the amastigote life stage

To understand how and why mitochondrial gene expression is regulated in *T. cruzi*, it was first necessary to understand expression in a mammalian as well as insect life stages. Therefore, we tested gene expression levels in the mitochondria in the amastigote life stage. Our previous work is the impetuous for these experiments, in which we showed that culture-derived trypomastigotes exhibit higher levels of many mitochondrial RNAs relative to log-stage epimastigotes. However, mRNA abundance is actually highest in the starved epimastigotes rather than in the trypomastigotes of a mixed culture (Shaw *et al*., 2016). Therefore, we wondered how transient the changes in gene expression upon starvation actually were. Are these changes retained once *T. cruzi* begins to replicate intracellularly?

We know from another study (Kalem *et al*., 2018) that expression of some nuclear-encoded ETC subunit mRNAs differ greatly between strains. Whether this reflects differences at the protein level is unknown, but the finding implies that mitochondrial RNA abundances of more than one strain should be tested to verify that potential findings are general rather than strain-specific. For this reason we performed these analyses in both the CL Brener strain used previously, and in the Sylvio X10 strain that was found to harbor substantial differences in nuclear gene expression (Kalem *et al*., 2018). Similarly, as *T. cruzi* will encounter and infect different cell types in different stages of human infection, our experiments included both fibroblasts (murine, 3T3) and cardiac cells (human, AC16; Davidson *et al*., 2005) as cultured host cells for *T. cruzi* infection.

The percentage of trypomastigotes in the mixed epimastigote/trypomastigote differentiation culture used to infect mammalian cells was determined for each strain. These numbers were 15 ± 7% (CL Brener) and 57± 21% (Sylvio X-10) in differentiation assays just prior to the infection experiments, although these rates varied extensively per differentiation. As column purification of culture-derived trypomastigotes to obtain 100% trypomastigote cultures could influence gene expression, we opted to infect directly with the mixed cultures, using a higher multiplicity of infection (MOI) than if we were using pure trypomastigotes. We took these percentages into account when deciding to use MOIs for CL Brener (200) that were higher than that of Sylvio X10 (50-100). Once mammalian cells were infected, both number of cells containing amastigotes, and number of amastigotes per infected mammalian cell were determined at 72 h post-infection (Fig. S1), when burst mammalian cells and cell-derived trypomastigotes were not yet observed in culture (not shown). The percent of fibroblasts infected was not different when infecting with CL Brener compared to Sylvio X10 strain, but this number was significantly different between cardiomyocyte infections (P = 0.0455). There was no significant difference between the number of amastigotes per cell between both *T. cruzi* strains and host cell lines, which provides evidence that all infections were in a similar phase of replication at this point.

Gene expression levels of mitochondrial RNAs were first compared between replicating amastigotes (collected at the same time that infection metrics were determined) and the *T. cruzi* culture originally used to infect these cells. As only twenty total protein coding and rRNA loci exist in the mitochondrion, and deep sequencing read populations of *T. cruzi* mitochondrial genomes contain relatively few that are fully canonically edited (Gerasimov *et al*., 2018), qRT-PCR is the best method to compare trypanosome mitochondrial RNA abundances for this study. We first examined the abundances of eight RNAs not affected by uridine insertion/deletion editing. Fig. 1A-B reveals that in comparison to the infecting culture, amastigotes of both *T. cruzi* strains exhibit a general decrease of abundance of mitochondrial RNAs unaffected by editing. The decreases in abundances that occurred in the replicating amastigote stage were remarkably similar between infections of murine fibroblasts and human cardiomyocytes, with only *ND2* (encoding a complex I subunit) and *MURF5* (maxicircle unidentified reading frame 5) differing significantly (P <0.05) in an unpaired t test. Decreases in abundances appeared to differ somewhat more between *T. cruzi* strains (compare Fig. 1A with Fig. 1B). Ribosomal rRNAs 9S and 12S and mRNAs *ND2* and *CO1* (encoding a complex IV subunit) exhibited the greatest decreases in abundance, with complex I subunit mRNAs *ND1, ND4, ND5* remaining largely unaffected.

**Figure 1.**
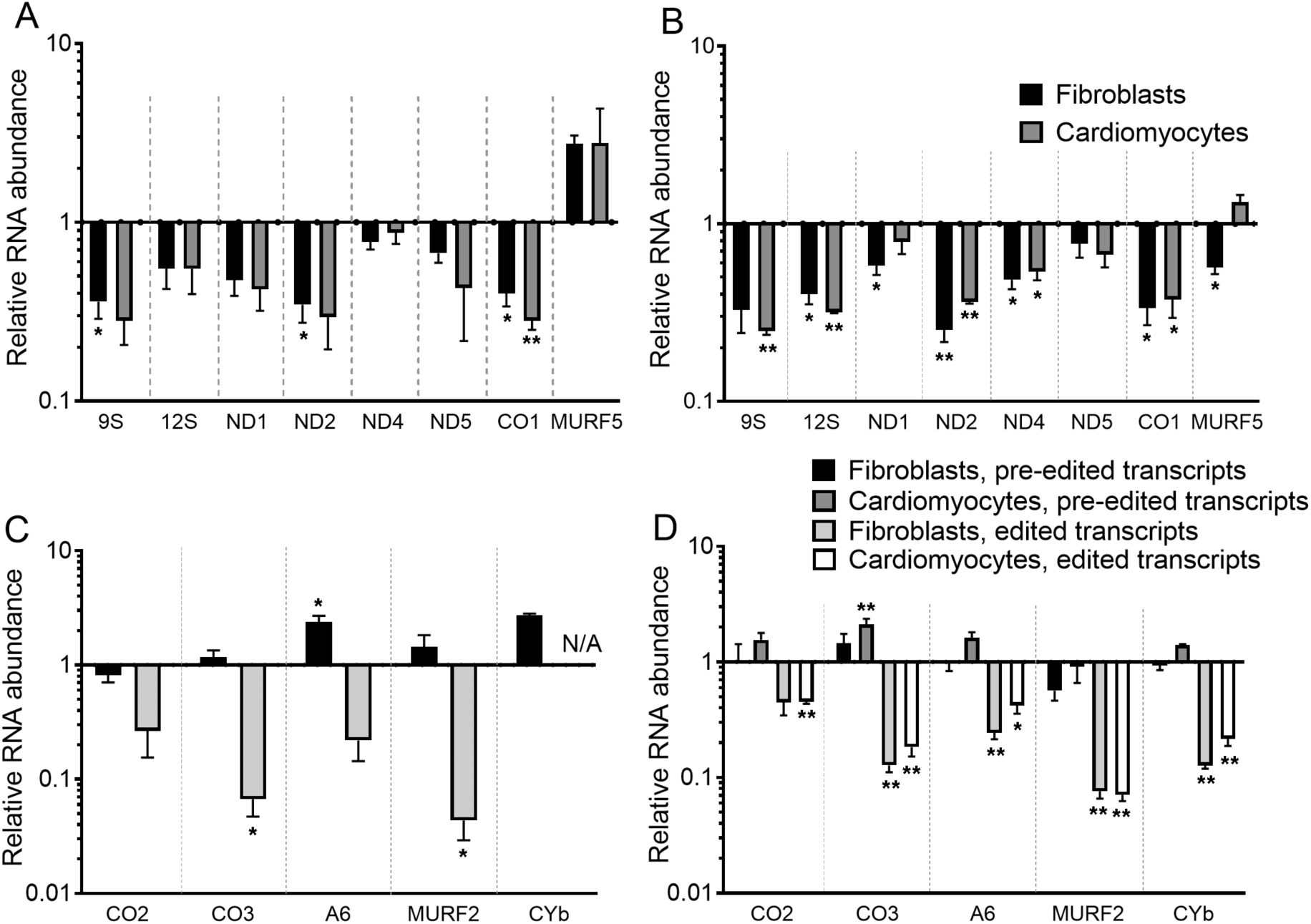
Mitochondrial RNA abundances in *Trypanosoma cruzi* intracellular amastigotes, ***relative to infecting T*. *cruzi culture abundances*. A. CL Brener** strain **never-edited** RNAs in amastigotes infecting fibroblasts and cardiomyocytes. **B**. Same as A, but with **Sylvio X10** strain infections. **C. CL Brener** strain **pre-edited and edited transcripts** in amastigotes infecting fibroblasts. Primer set for edited *CYb* did not amplify acceptably in either life stage. **D. Sylvio X10** strain **pre-edited and edited transcripts** in amastigotes infecting fibroblasts and cardiomyocytes. Value of 1 on *y* axis means no difference in the RNA abundance relative to infecting cultures. ** (P < 0.01) and * (P <0.05) indicate results of a ratio paired t test of amastigote mRNA level in comparison to that of infecting culture. Data points represent the mean of three biological replicate infection assays per cell line. Error bars represent standard error of the mean. Normalization was to the mean abundance of *TERT* and *PFR*.

The mature transcripts for the remaining mitochondrial mRNAs are generated after completion of uridine insertion/deletion RNA editing, so relative amastigote RNA abundances of these mRNAs, before and after editing, was examined next. We examined the same transcripts that we had tracked abundance of during starvation and culture differentiation (Shaw *et al*., 2016): three mRNAs edited in a small, defined region, and two mRNAs edited throughout their entire coding region (extensively-edited; Fig. 1C-D). We saw a different pattern in relative abundances with these mRNAs. While the pre-edited versions of mRNAs remained consistent or even increased up to two-fold in abundance, edited mRNA abundances were always lower, often dramatically so (ranging from two to twenty-fold less), in the dividing amastigotes. While analysis was only performed on fibroblast infections of CL Brener, comparison of abundances in strain Sylvio X10 amastigotes in fibroblast and cardiomyocyte infections reveal nearly identical values with the exception of *CYb* (encoding a complex III subunit) levels, where depletion levels of pre-edited and edited forms differed significantly, P <0.05, in an unpaired t test, again suggesting little impact of the identity of the infecting cell line. Overall, abundance profiles of amastigote edited mRNAs suggest that cell cycle-specific control of abundance of mature mitochondrial mRNAs is performed at the level of editing when possible.

Comparison of Fig. 1 relative abundance patterns and those detected previously between replicating, fed epimastigotes and starved, unreplicating cells of mixed morphology (Shaw *et al*., 2016) suggested that there may be a low, consistent level of mitochondrial mRNA expression of *T. cruzi* as it replicates, regardless of whether it is in amastigote or epimastigote life stage. To determine if this is true, we compared mRNA abundance values obtained in amastigotes with those of log-phase epimastigotes (Fig. 2). Relative to exponentially growing cells, most never-edited transcripts were statistically lower in abundance in amastigotes, contrary to our expectations. Abundances of several transcripts remained unchanged in the pre-edited and fully edited mRNA sequences, and in *CO3* (encoding a complex IV subunit) there were significantly higher abundances of the pre-edited transcript. Edited *CO3, MURF2* (maxicircle unidentified reading frame 2) and *CYb* were dramatically reduced in amastigotes. Overall, these results show that there is a general trend of lower translatable mtRNA transcript abundances in Sylvio X10 amastigotes when compared to insect-stage epimastigotes, revealing a difference in the mitochondrial transcriptome between these two replicating life stages. However, the analysis also showed that the magnitude of difference is lower than when comparing either replicating stage to non-replicating *T. cruzi*. While the implications of the difference in *MURF2* cannot be hypothesized as its function is not known, the lower abundance of many ETC subunit transcripts in amastigotes could indicate differences in the roles of ETC complex III or IV between the two replicating life stages.

**Figure 2.**
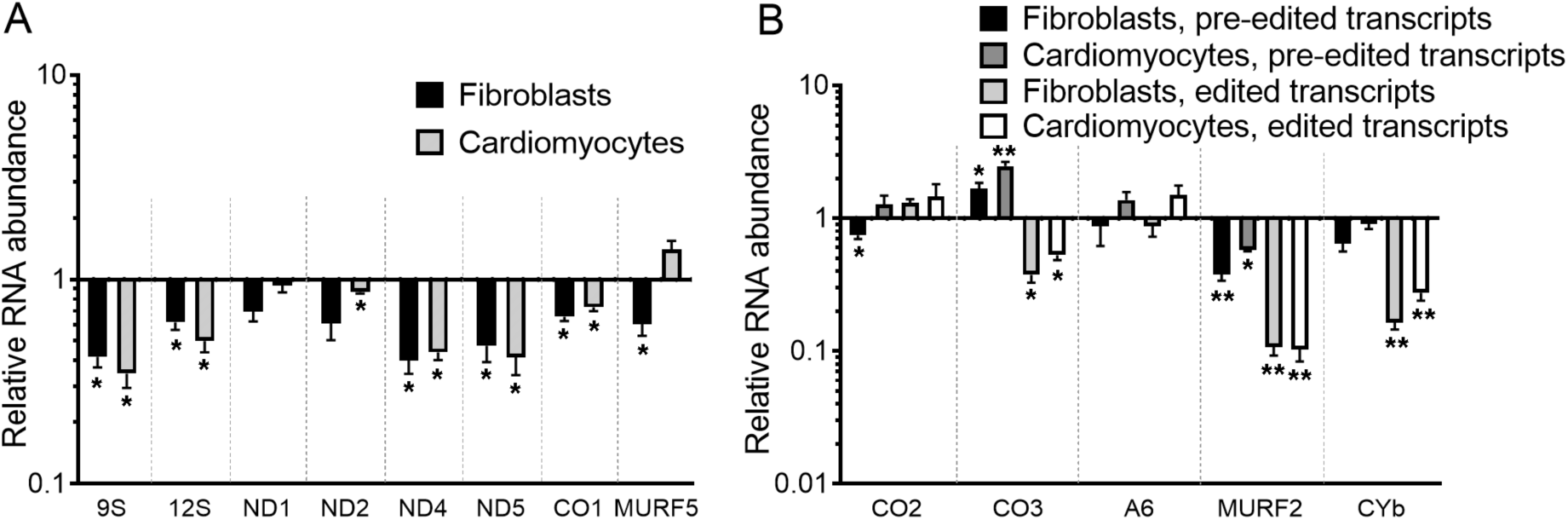
Mitochondrial RNA abundances in *Trypanosoma cruzi* Sylvio X10 strain amastigotes, ***relative to exponentially growing culture epimastigotes*. A. Never-edited** mitochondrial RNAs **B. Pre-edited and edited** mitochondrial transcripts. Value of 1 on *y* axis means no difference in the RNA abundance relative to that of fed epimastigotes. ** (P < 0.01), * (P < 0.05) indicate results of a ratio paired t test in comparison to exponentially growing cells. Data points represent the mean of three biological replicate infection assays each in fibroblasts and cardiomyocytes. Error bars represent standard error of the mean. Normalization was to the mean abundance of *TERT* and *PFR*.

Mitochondrial RNA abundance comparisons thus far have shown trends consistent between *T. cruzi* strains. To establish that this is also true of comparisons between starved and unstarved epimastigotes, we repeated experiments of acute starvation by incubation in TAU for 48 h. as performed previously using strain CL Brener (Shaw *et al*., 2016) on a limited number of transcripts and found similar patterns although abundance increases upon starvation were more robust in general in Sylvio X10 (Fig. 3A). Finally, we were interested to know whether mitochondrial mRNA levels differ between these strains in a single life stage or condition, either the infective culture (containing trypomastigotes) or the amastigote life stage. Most mRNAs examined were approximately equally expressed in both strains relative to the two nuclear-encoded normalization genes. Exceptions largely consisted of *ND2* and edited mRNAs that were always at higher levels in the Sylvio X10 culture. This may indicate a higher level of editing in Sylvio X10 in general, although other explanations should not be ruled out. *MURF5* and a few pre-edited mRNAs were also more abundant in Sylvio X10, but only in the mixed infective culture (Fig. 3B). From Fig. 3 experiments we conclude that life stage and starvation-triggered variation of *T. cruzi* mitochondrial gene expression is a phenomenon likely conserved among strains, and that absolute abundance of mitochondrial mRNAs is largely conserved, with exceptions.

**Figure 3.**
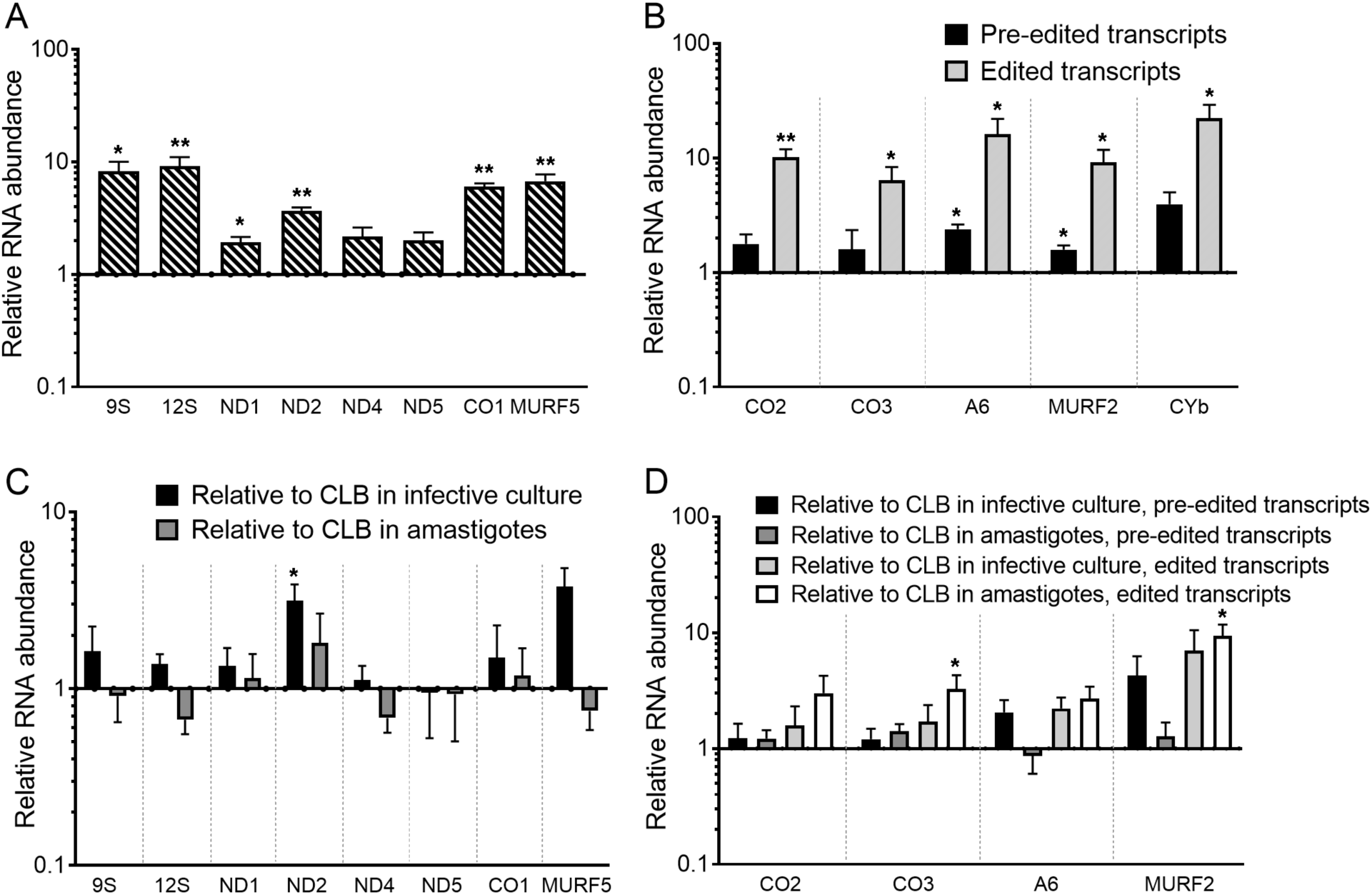
Mitochondrial RNA abundances in *Trypanosoma cruzi* Sylvio X10 strain. **A**. Abundance of **never edited** RNAs in TAU-starved epimastigotes, relative to exponentially growing cells. **B**. Abundance of **pre-edited and edited** transcripts in TAU-starved epimastigotes, relative to exponentially growing cells. ** (P < 0.01), * (P < 0.05) in A. and B. indicate results of a ratio paired t test in comparison to exponentially growing cells. **C**. Abundance of **never edited** RNAs in Sylvio X10 strain infective culture and amastigotes, relative to levels in the same CL Brener strain (CLB) life stage (performed in fibroblasts) **D**. Same as C. except abundance of **pre-edited and edited** transcripts. Asterisks (*, P < 0.05) in C. and D. indicate results of a multiple t tests corrected for multiple comparisons by the Holm-Sidak method. Value of 1 on *y* axis means no difference in the relative RNA abundance. Data points represent mean of three biological replicates. Error bars represent standard error of the mean. Normalization was to the mean abundance of *TERT* and *PFR*.

In experiments shown in Figs. 1–3, the two maxicircle unidentified reading frames *MURF5* and *MURF2* (named for the large circular component of the trypanosome mitochondrial genome), were often exhibiting patterns differing from the rest of the transcripts. For example, *MURF5* was upregulated rather than decreased in CL Brener strain in intracellular amastigotes relative to the infecting culture. *MURF2*, like *MURF5*, seemed to be more abundantly expressed in Sylvio X10 compared to the CL Brener strain. Previous investigation of the maxicircle has shown that possibly these loci may not be true open reading frames in *T. cruzi* as their DNA sequences in some strains contain internal stop codons or frameshifts (Ruvalcaba-Trejo and Sturm, 2011). Therefore, we may be observing noise due to variable processing of the surrounding transcripts that likely are polycistronically transcribed, for which precursor mRNAs would contain *MURF* qRT-PCR target regions. More interesting are differences in abundance between strains or modulation across the life cycle for mRNAs encoding ETC complex I ETC components, but our understanding of *T. cruzi* complex I is also very limited (Carranza *et al*., 2009).

### rRNAs that increase in abundance upon starvation appear to be incorporated into ribosomes

It is a mystery why some mitochondrial RNAs would exhibit abundance spikes during starvation and in the trypomastigote stage that is nonreplicating. Theoretically these conditions could reduce rather than increase the need for these RNAs. This is particularly true as most mitochondrial mRNAs encode ETC subunits, and two nuclear-encoded ETC subunits were not found to increase in abundance during starvation ((Shaw *et al*., 2016); antibodies cannot be generated to subunits encoded in the mitochondrial genome). To explore this further we asked whether the rRNAs that were increased in starved epimastigotes were incorporated into ribosomes or instead accumulated as non-associated RNAs. Under conditions of 48 h of severe starvation that leads to a 6-fold increase in 12S rRNA in strain CL Brener (Shaw *et al*., 2016), we hypothesized that unincorporated rRNAs would be observable in low molecular weight fractions of cell lysate separated on a sucrose gradient. In contrast, if polysome profiles were the same prior and post-starvation, with no prominent bump of rRNA in low molecular weight fractions, it suggests that the extra rRNAs are being incorporated into ribosomes or ribosomal subunits. This is what we observed. Fig. 4 shows normalized signal abundances for mitochondrial rRNA (detected by RNA blot with radiolabeled DNA oligomer probes) summed in fractions containing non-associated nucleic acid and proteins, individual ribosome subunits, ribosomes, and polysomes. No statistical differences are observed between fractionation of lysate from replicating fed epimastigotes and starved cells, and no differences of a magnitude that we would expect given a six-fold increase in rRNA expression.

**Figure 4.**
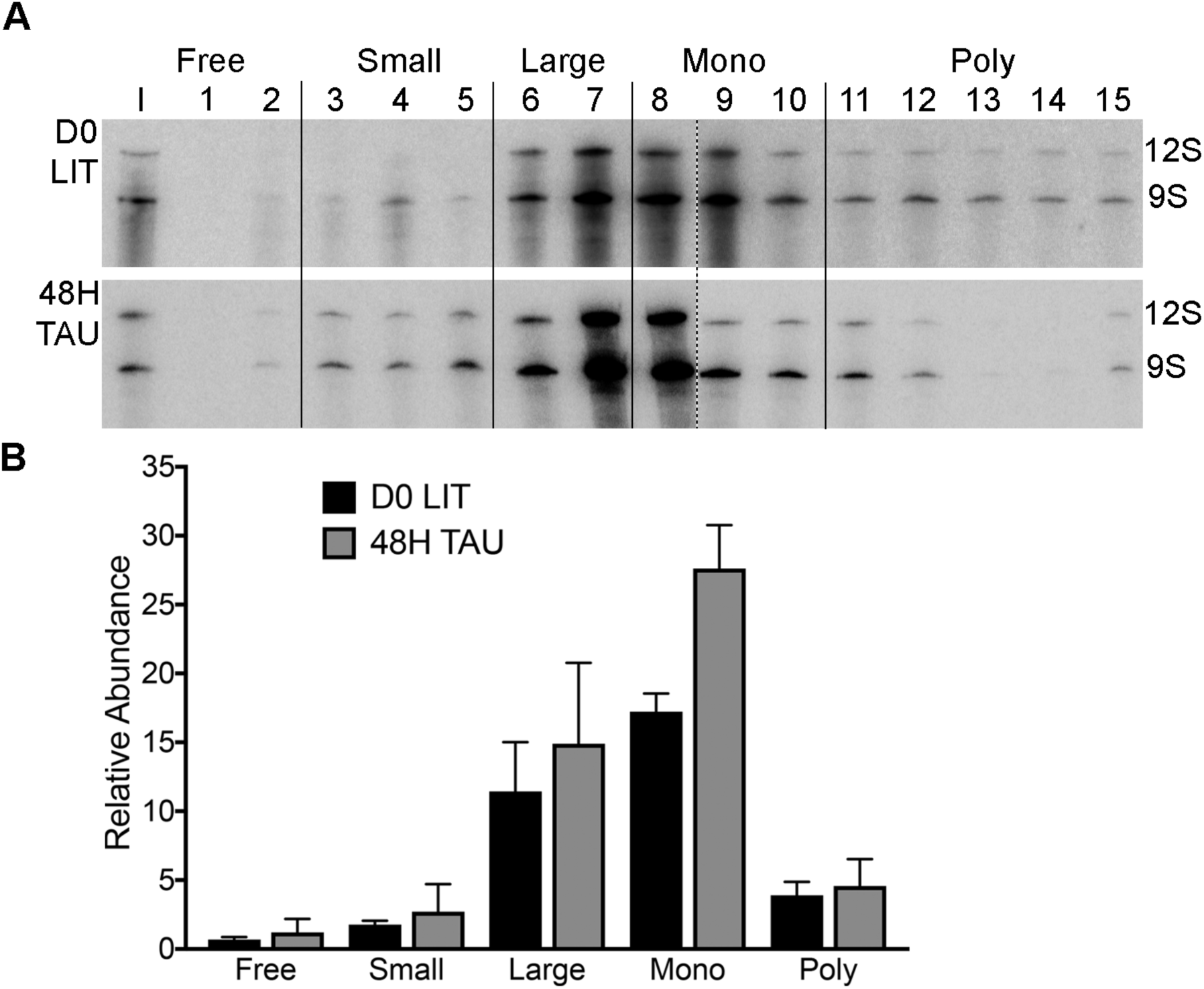
Mitochondrial ribosomal RNA contained in specific segments of *Trypanosoma cruzi* mitochondrial polysome profiles. Free, unassociated rRNA; Small, small subunit-associated rRNA; Large, large subunit-associated rRNA; Mono, monosome; Poly, polysome. Discrete polysome segments were determined in comparison to gradient positions of bacterial rRNA migration. **A**. Representative RNA blot probed with 12S and 9S oligonucleotide probes during exponential growth in LIT medium (D0 LIT) and starvation in TAU medium (48H TAU). Dotted line indicates boundary between 2 separate blots that contain the sum of the gradient fractions. **B**. 12S and 9S densitometry signals were obtained from RNA blots from each gradient fraction and normalized to that of a total input lane for that experiment. Data represents the mean of three biological replicate experiments for each condition. Error bars represent standard error of the mean.

### *Multiple aspects of* T. cruzi *mitochondrial respiratory output are decreased upon starvation*

If extra ribosomes are present in the starved epimastigotes and likely in trypomastigotes, they may be either actively being used or sequestered. Since testing either mitochondrial-encoded protein abundances or translation of the mitochondrial genome is a technical hurdle, we instead sought to determine whether we would see an increase in respiration that would suggest higher levels of ETC complexes or subunits. Although increases in respiration in non-dividing cells is counter-intuitive, such a phenomenon may occur when substrate availability issues (loss of environmental glucose) profoundly alter *T. cruzi* metabolism (Barisón *et al*., 2017). We analyzed mitochondrial respiration of CL Brener strain cells harvested in exponential and stationary growth phases in normal medium (D0 LIT and D8 LIT, respectively), and abrupt, severe starvation (48H TAU). Overall respiratory output (Fig. 5) was found to decrease in the starvation conditions, with the largest decreases attributed to abrupt starvation. ATP-coupled respiration and spare respiratory capacity appeared particularly affected (Fig. 5B and D). In this respect, the cells appear to function as is typical for cells generally under conditions of growth cessation due to nutrient unavailability. Therefore, in contrast to the fact that culture starvation yields more mRNAs encoding ETC components during starvation, the overall output of this complexes decreases during the same starvation conditions. Possibly, extra ribosomes generated during this time are not actively translating their associated mRNAs. Indeed, although not statistically significant, there does appear to be a trend toward an increase in monosomes relative to polysomes in profiles derived from starved cells (Fig. 4). Finally, the paradoxical increase in mRNA abundances coupled with less respiration in starved epimastigote cultures suggests that we should not necessarily equate low abundances of mitochondrial RNAs in intracellular amastigotes with fewer ETC complexes or activity. In fact, amastigotes respire (Li *et al*., 2016).

**Figure 5.**
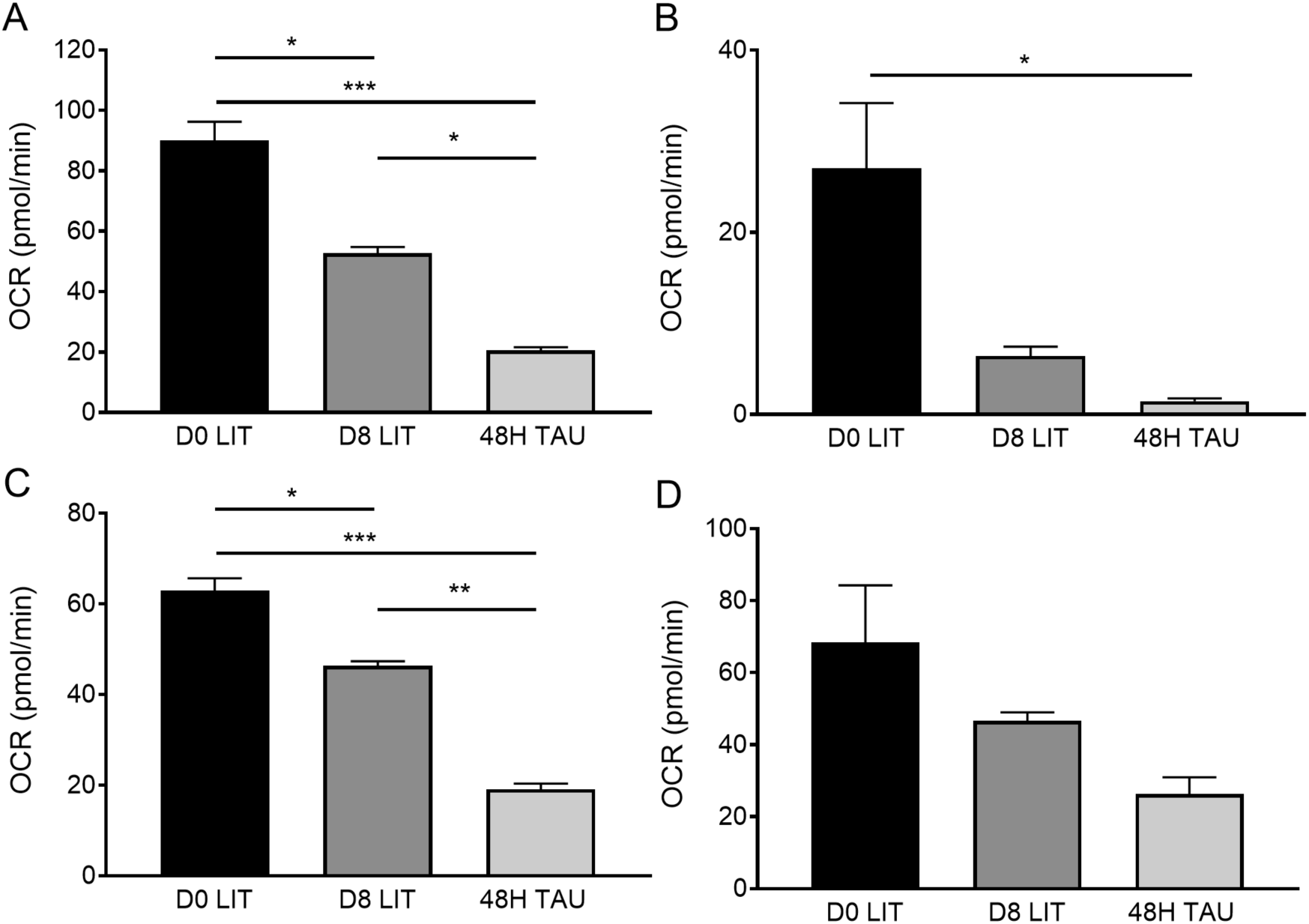
Mitochondrial electron transport chain (ETC) utilization in *Trypanosoma cruzi* CL Brener. **A**. Basal respiration measured prior to ETC inhibitor addition (ANOVA P<0.0001). **B**. ATP-coupled respiration (ANOVA P=0.0121). **C**. Proton leak (ANOVA P=0.0179). **D**. Spare respiratory capacity (ANOVA P<0.0001). Oxygen consumption rate (OCR) measurements are normalized to relative SYBR Green 1 fluorescence levels in each well. Data show the means of four biological replicates of six technical replicates each. Error bars represent standard error of the mean. * represents p *≤* 0.05, ** represents p *≤* 0.001, and *** represents p *≤* 0.0001

### Tc*RND overexpression results in reduction of extensively edited transcripts*

Variations in mitochondrial RNA abundance and editing across the life cycle and during starvation are dramatic. However, we cannot show that they result in logically expected functional changes for the cell, leading us to approach the paradox from another direction. We employed a genetic approach to depress editing, and evaluated the effects on cultured CL Brener fed and differentiating epimastigotes. Elimination of the critical components of the editing machinery would likely be toxic for *T. cruzi*, and inducible methods for knock-out or reduction of these proteins is not yet available. However, when inducibly overexpressed, *T. brucei Tb*RND, a uridylate-specific exoribonuclease that appears specific to gRNAs, severely depletes levels of extensively edited mitochondrial mRNAs. As a system for inducible expression in *T. cruzi* does exist, we attempted this approach, targeting the *Tb*RND *T. cruzi* homologue, *Tc*RND. Our plan was to use a tetracycline (tet) based system to induce ectopic, untagged *Tc*RND expression in addition to continued native *Tc*RND expression, expecting that the cumulative expression would cause depletion in extensively-edited mitochondrial transcripts as in *T. brucei*.

A Gateway® cloning-compatible (Alonso *et al*., 2014) pTcINDEX system was used to express *Tc*RND integrated into the transcriptionally silent rRNA spacer region in *T. cruzi* constitutively expressing T7 polymerase and tet repressor episomally (Taylor and Kelly, 2006). Experiments were performed in two backgrounds. In the first, T7 polymerase and tet repressor were supplied by transfection of the *T. brucei* plasmid pLEW13 (Wirtz *et al*., 1999); in the second, a pLEW13 derivative in which T7 polymerase and tet repressor expression was driven by the *T. cruzi* rRNA promoter (p*Tc*rRNA-T7Tet; (Taylor and Kelly, 2006)) was used. Following transfection and selection of resistant cells, expression of *Tc*RND mRNA was determined by reverse transcription and quantitative PCR (qRT-PCR).

Fig. 6A contains results from three cell lines. Two were obtained from the pLEW13 background from separate transfections, and one was derived from the p*Tc*rRNA-T7Tet (T7) parent line. In all cases, *Tc*RND-transfected cells had much higher levels of *Tc*RND mRNA relative to both the parent line and untransfected parasites. Expression of the extra copy of *Tc*RND was always leaky; in the case of the pLEW13-derived *Tc*RND transfected cell lines, 13RND1 and 13RND2, five to tenfold increased expression was observed even without *tet* induction, and *tet* induction for 48 h resulted in an additional five to tenfold induction. Leaky expression is more profound in the transfected *Tc*RND cells from the T7 cell line, T7RND. Here, hundredfold overexpression of *Tc*RND is observed regardless of induction. Levels of *Tc*RND upon *tet* induction were consistent for 8 days, the longest time tested (not shown).

**Figure 6.**
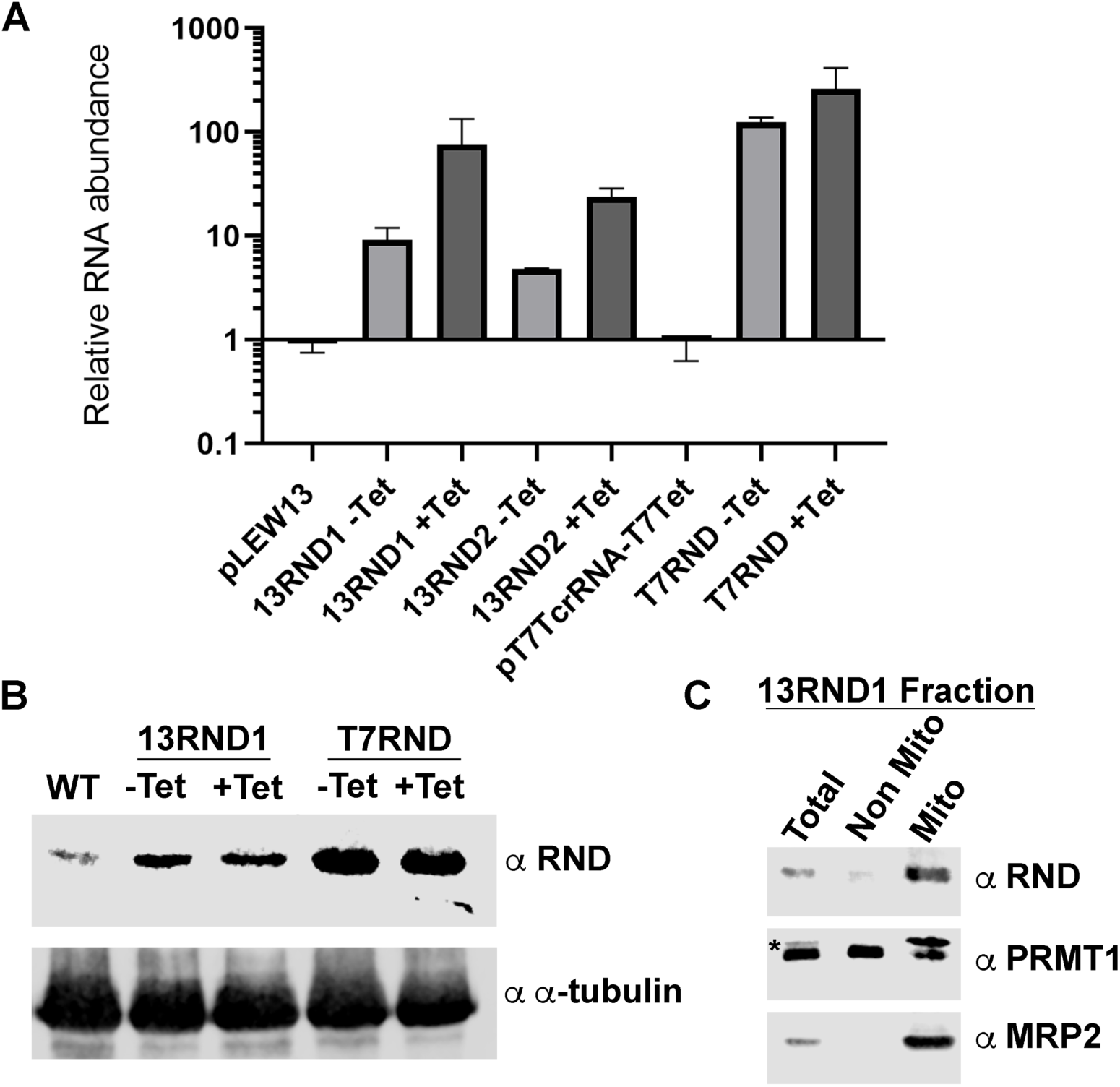
**A**. Abundance of *Tc*RND transcript in *Trypanosoma cruzi* strain CL Brener transfected with pLEW13 (parent pLEW13), p*Tc*rRNA-T7Tet (parent p*Tc*rRNA-T7Tet), and 48 h induced (+Tet) and uninduced (-Tet) 13RND1, 13RND2, and T7RND parasite cell lines ***relative to untransfected CL Brener strain*** (wild-type cells). Value of 1 on *y* axis means no difference in Tc*RND* abundance relative to that of untransfected CL Brener strain. Data show the means of two biological replicates (two technical replicates each). Error bars represent standard error of the mean. Normalization was to the mean abundance of *TERT* and *PFR*. **B**. Immunoblot analysis of expression of *Tc*RND in indicated parasite cell lines. Tubulin is the loading control. **C**. Immunoblot analysis of subcellular fractionation of *Tc*RND in cell line 13RND1. Loaded are total lysate (Total), non-mitochondrial fraction consisting of the supernatant following centrifugation of crude mitochondrial vesicle pellet (Non Mito; clarified lysate), and mitochondrial vesicle fraction protein (Mito). PRMT1 is a non-mitochondrial protein and MRP2 is a mitochondrial protein used as controls. * indicates non-specific band recognized in *T. cruzi* by *T. brucei* PRMT1 antibody. In both B. and C., signal recognized by RND antibody derives from both native *Tc*RND and that expressed from an intergenic rRNA locus. Tet induction is for 48 h.

We next verified that this overexpression persisted at the protein level. Serum containing *Tb*RND antibodies (Zimmer *et al*., 2011) nonspecifically identified many *T. cruzi* proteins. Antibodies were purified from the serum by their ability to bind recombinant *Tc*RND, as *Tb*RND and *Tc*RND amino acid sequences are >77% identical and ~85% similar. The newly-purified RND antibody still recognizes *Tb*RND very specifically (Fig. S2). Additionally, it recognizes *Tc*RND from total cell lysate of the same size as the *T. brucei* protein (Fig. 6B). At the protein level, *Tc*RND overexpression appears robust but fairly similar between induced and uninduced cells in transfected cell lines relative to the untransfected parental cells. However, *Tc*RND expression is higher in the T7RND line than in either 13RND line. We also determined that overexpressed *Tc*RND was correctly localized in 13RND1 (Fig. 6C). In summary, we have generated multiple cell lines that overexpress TcRND but are marginally or not at all inducible.

The effect of TcRND on growth in was examined in these cell lines. In *T. brucei*, overexpression of TbRND results in severe growth inhibition that is dependent on its enzymatic activity (Zimmer *et al*., 2011). Surprisingly, we saw that the cell lines that overexpressed TcRND grew at normal or near-normal rates (Fig. 7A). However, it does appear that maximum concentration of cells reached as the culture enters stationary phase may be somewhat compromised when *Tc*RND is overexpressed in the 13RND1 cell line. As the *Tc*RND levels were so high yet growth unaffected in the T7RND parasites, they potentially have compensatory mutations that developed prior to most of the testing. Further experiments demonstrate it to be the least reliable of the three cell lines.

**Figure 7.**
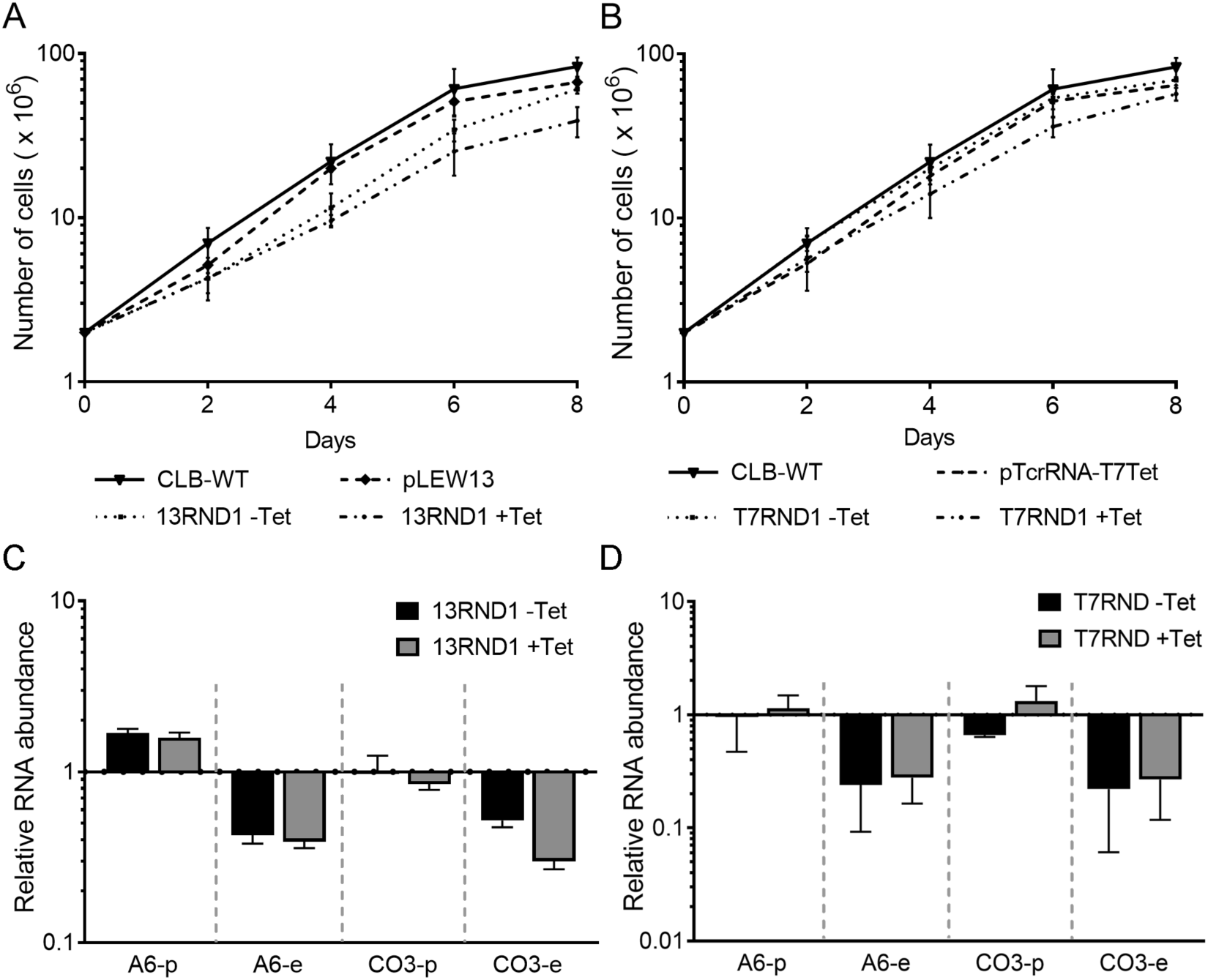
Growth of *Tc*RND overexpressing *Trypanosoma cruzi* cell lines. **A**. Growth curve of 13RND1 (48 h induced and uninduced) compared to the background (pLEW13-expressing) and wild-type, untransfected strain CL Brener. **B**. Growth curve of T7RND (48 h induced and uninduced) compared to the background (p*Tc*rRNA-T7Tet-expressing) and wild-type, untransfected strain CL Brener. Data show means of three replicates of each cell line (wild-type CL Brener shown twice, once in each panel for comparison). Error bars represent the standard error of the mean (SEM). One-way analysis of variance (ANOVA) indicated that cell concentrations were not statistically different from one another at all timepoints (p>0.05). **C**. Abundance of pre-edited and edited mitochondrial transcripts in uninduced (-Tet) and 48 h induced (+Tet) *Tc*RND overexpressing parasites (p13RND1), relative to CLB-WT. Value of 1 in *y* axis means no difference in RNA abundance relative to untransfected CL Brener strain. Data show the means of three biological replicates (two technical replicates each). Error bars represent standard error of the mean. Normalization was to the mean abundance of *TERT* and *PFR*. **D**. Like C., except analyzing levels of mitochondrial transcripts in the T7RND parasite cell line, and two biological replicates were analyzed.

We were concerned that the relatively normal growth rates indicated non-functionality of the *Tc*RND expressed from the rRNA intergenic locus. However, mitochondrial mRNA abundance analysis suggests that the overexpressed *Tc*RND is active (Fig. 7B). In *T. brucei, Tb*RND overexpression results in the depletion of the edited but not the pre-edited species of extensively edited mRNAs such as *CO3* and *A6* (Zimmer *et al*., 2011). We analyzed these mRNAs in the 13RND1 line epimastigotes and found results similar to that of the *T. brucei* overexpressor. We saw approximately two-fold to four-fold depletion of exclusively the edited forms of these mRNAs. This indicates that this role of *Tc*RND in *T. cruzi* likely mirrors that of the *T. brucei* homologue. Similar results were observed with the T7RND line (Fig. 7C) and the second pLEW13-derived line (not shown). Therefore, *Tc*RND overexpression results in disruption of editing, at least of extensively-edited transcripts. Its negligible effect on epimastigote growth, while surprising, makes the fact that we are unable to control the overexpression less of a problem in these experiments. It is also the reason that we did not more aggressively pursue alterations of selective pressures and tet levels and other factors that may have resulted in our obtaining an inducible *Tc*RND overexpressor with tighter control, as is theoretically possible to achieve (Tavernelli *et al*., 2019).

### TcRND overexpression affects epimastigote physiology and differentiation

With our *Tc*RND ovexpressing cells we could ask whether a compromised ability to generate edited (and thus translatable) mRNAs impacts *T. cruzi*’s ability to differentiate to the trypomastigote form in culture. We performed culture differentiation using our slow starvation and low fetal bovine serum protocol (Shaw *et al*., 2016) in untransfected parasites, parent pLEW13, and uninduced and induced 13RND1. The results (Fig. 8A) revealed a dramatic reduction of differentiation to the trypomastigote life stage for 13RND1. A non-significant reduction in percentage of cells that are trypomastigote was observed in the pLEW13 parent line relative to untransfected cells, but when the cells overexpressed *Tc*RND, the percentage of cells that had successfully differentiated was reduced by over half, from 27.3±2.1% to 10.4±1.3% in the induced 13RND1 cell line. We also analyzed the effect of *Tc*RND overexpression in the traditional culture differentiation model, with consistent results (Fig. S3). Therefore, the compromised ability to execute mitochondrial mRNA editing that appears fairly innocuous in fed epimastigotes (very little impact on growth), profoundly affects *T. cruzi*’s ability to differentiate.

**Figure 8.**
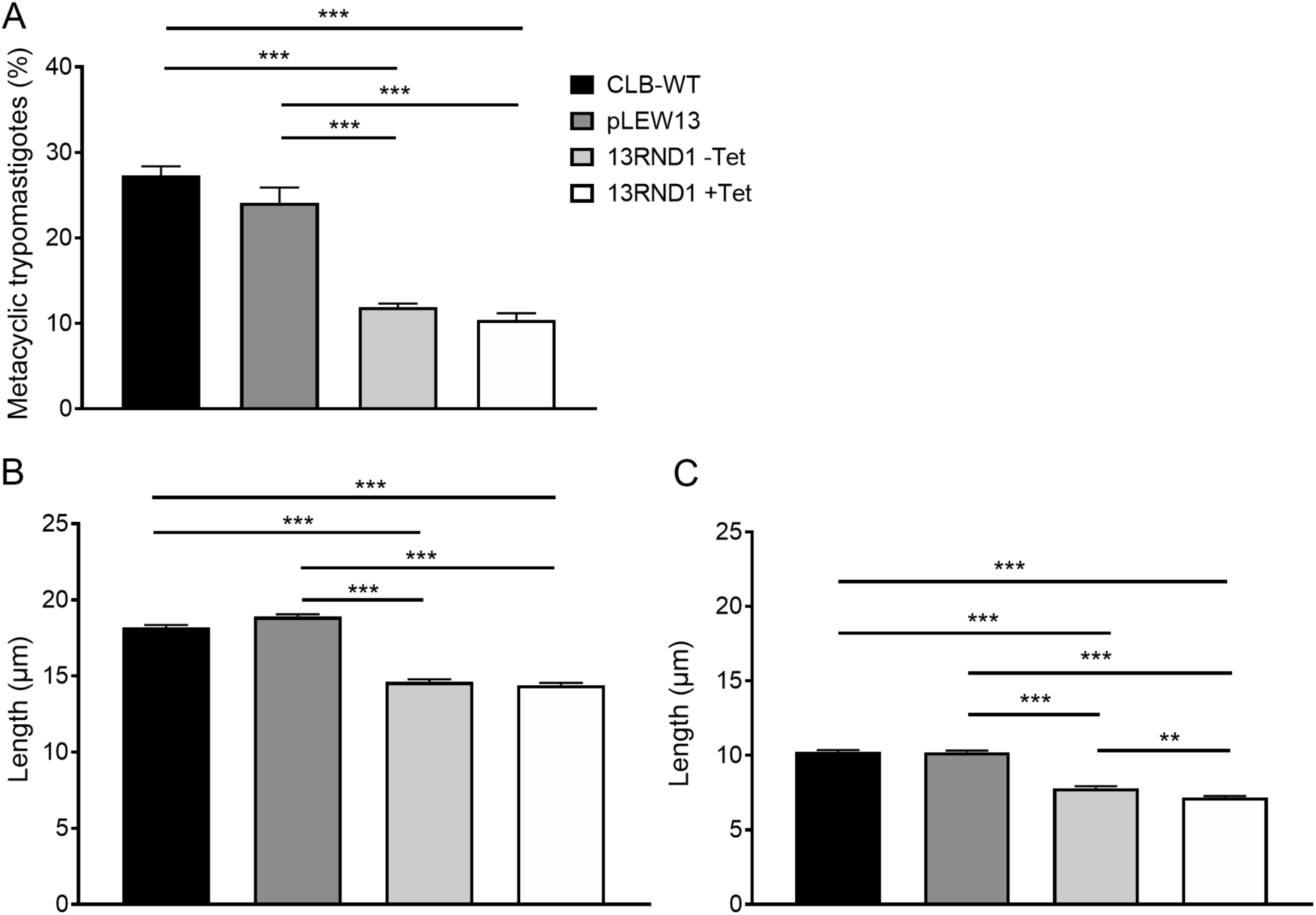
Phenotypic changes associated with *Tc*RND overexpression in *Trypanosoma cruzi*. **A**. Percentage of metacyclic trypomastigotes of wild-type strain CL Brener (CLB-WT), pLEW13-expressing parent strain (pLEW13), induced (+Tet) and uninduced (-Tet) *Tc*RND overexpressing parasites (13RND1) in mixed, differentiating cultures after 8 days in restricted medium (ANOVA P<0.0001). Cultures were induced for 48 h prior to initiation of differentiation. **B**. Total average length of CLB-WT, pLEW13, induced (+Tet) and uninduced (-Tet) 13RND1 epimastigotes in exponential growth (ANOVA P<0.0001). Cultures were induced or not and allowed to grow for 48 h prior to analysis. **C**. Free flagellum length of CLB-WT, pLEW13, induced (+Tet) and uninduced (-Tet) 13RND1 as in B. (ANOVA P <0.0001). At least 100 cells were counted or measured from each replicate. For all experiments, data show mean of three biological replicates. Error bars represent standard error of the mean. * p *≤* 0.05, ** p *≤* 0.001, and *** p *≤* 0.0001.

Differentiation is intricately linked to other cellular events that precede it. For instance, cell and flagellar lengthening of epimastigotes is observed as cells exit exponential growth and enter stationary phase as a result of nutrient depletion (Hernandez *et al*., 2012). It has also been determined that the loss of glucose in culture is the trigger for flagellar elongation (Tyler and Engman, 2000). Lengthening of cells and flagella is hypothesized to be “preadaptive” for metacyclogenesis (Tyler and Engman, 2000; Hernandez *et al*., 2012), although the connection remains obscure and the natural isolates contain a mix of insect life stages and are extremely morphologically varied within and between isolates (Kollien and Schaub, 1998; Silva *et al*., 2019).

While working with our *Tc*RND-transfected cell lines, we noticed that they appeared shorter than their wild-type counterparts, even in log-stage epimastigote culture. We therefore measured the cells for a quantitative assessment and comparison. Indeed, epimastigotes that overexpress *Tc*RND are shorter overall (Fig. 8B). In order to determine whether the body or the flagella was contributing to the length difference, both body and flagella were measured, and the greatest difference between normal and *Tc*RND overexpressing cells was flagellar length (Fig. 8C). In this parameter, a statistical difference was even found between uninduced and induced 13RND1. Similar differences were found in the other two cells lines (not shown). If this cell size phenomenon is related to the *T. cruzi*-specific insect stage differentiation event, we would expect that *Tb*RND overexpression in *T. brucei* does not result in any cell length alterations, and we found that it does not (Fig. S4). In summary, a length attenuation phenotype and a differentiation phenotype are features of mitochondrial *Tc*RND overexpression, and causative mechanisms for these phenotypes are likely to involve edited mitochondrial mRNA abundances.

## Discussion

This study provided knowledge of changes to mitochondrial gene expression throughout the *T. cruzi* life cycle, and then focused on the functional relevance of the spike in abundances of certain mRNAs during starvation and differentiation in culture. We have generated a model of how these findings fit together that also addresses the necessity for coordination of mitochondrial function with the rest of the parasite cell in the face of life cycle or environmental changes that we discuss here.

First, however, some experimental results must be addressed further. One is that continuously-growing cells appear fairly immune to decreased levels of mature mRNAs encoding ETC subunits caused by *Tc*RND overexpression. This result could indicate that the stability of protein product is very high, that efficiency of mitochondrial translation is primed to compensate for lower levels of mature mRNAs, or that the ETC is not used to capacity in normal parasite axenic culture that is rich in many energy sources. It is also apparent that *Tc*RND has a limited ability to impact editing, as even under conditions of high overexpression (Fig. 6) mature mRNAs are not depleted more than about four-fold. This may be due to the fact that *Tc*RND must exert its effects on a substrate (gRNAs), and mild overexpression is sufficient to completely target the population of gRNAs that are vulnerable to *Tc*RND activity.

An additional observation was made during the differentiation experiments that may be valuable to investigate in the future. Flasks differentiated by the method of Contreras *et al*. (1985) were examined every other day under the microscope (This examination was not performed in our normal differentiation protocol that calls for 8 days of undisturbed incubation). Parasites with normal *Tc*RND levels were mostly adherent to the flask surface during this protocol as expected, while p13RND1 parasites were not. Multiple sources cite adherence by the flagella as a component of metacyclogenesis (e.g. Bonaldo *et al*., 1988; Figueiredo *et al*., 2000; Hamedi *et al*., 2015). Further, starved cells in culture with lengthened morphology, including longer flagella, are more adherent (Tyler and Engman, 2000). Therefore, shorter flagella in 13RND1 could conceivably be directly responsible for its failure to differentiate at normal levels, although this conclusion remains speculative.

### Model connecting mitochondrial gene expression and differentiation

Almost nothing is known about coordination of mitochondrial and cellular processes in protists. In better-studied eukaryotic systems, it is accepted that mitochondria can signal to the rest of the cell aspects of their functional state and exert a cellular response. However, the sum of the mechanisms underlying the mitochondrial retrograde signaling and nuclear and cellular responses is still being established. Importantly, various species of noncoding RNAs originating from the nuclear genome have been identified within the mitochondrion, and although these findings are controversial due to the technical challenges of precise subcellular fractionation (Vendramin *et al*., 2017), evidence continues to emerge (Sirey *et al*., 2019). Transport of RNA out of the mitochondrion is also unresolved although hypothesized (Vendramin *et al*., 2017). Therefore, it is at least plausible that an RNA species from the mitochondrion might act as a mitochondrial retrograde signaler.

One of our major conclusions is that replicating life stages assayed in axenic or mammalian cell culture have fairly consistent levels of mature mRNA abundances that are lower than that of cultures of mixed infectious cultures that are not replicating. In Fig. 9, we propose that much of the trypomastigote ETC is lost or inactivated. These trypomastigotes therefore must possess a stockpile of ribosomes and mitochondrial mRNAs in order to regenerate ETC complexes after their loss/inactivation in the transitional period. It then is logical that epimastigotes would begin to accumulate these materials even prior to differentiation to trypomastigotes, and the differentiation will be blocked unless these levels are achieved. In our work, these checkpoints are manifest as reductions in flagellar extension and differentiation in the RND overexpressing parasites. Whether an mRNA would actually be transported out of the mitochondrion, or else its increasing presence would generate another signal that would be transmitted out we are currently unable to speculate on. However, the presence of retrograde transport of RNA (tRNA) from cytoplasm to nucleus has been already been established in *T. brucei* (Kessler *et al*., 2018).

**Figure 9.**
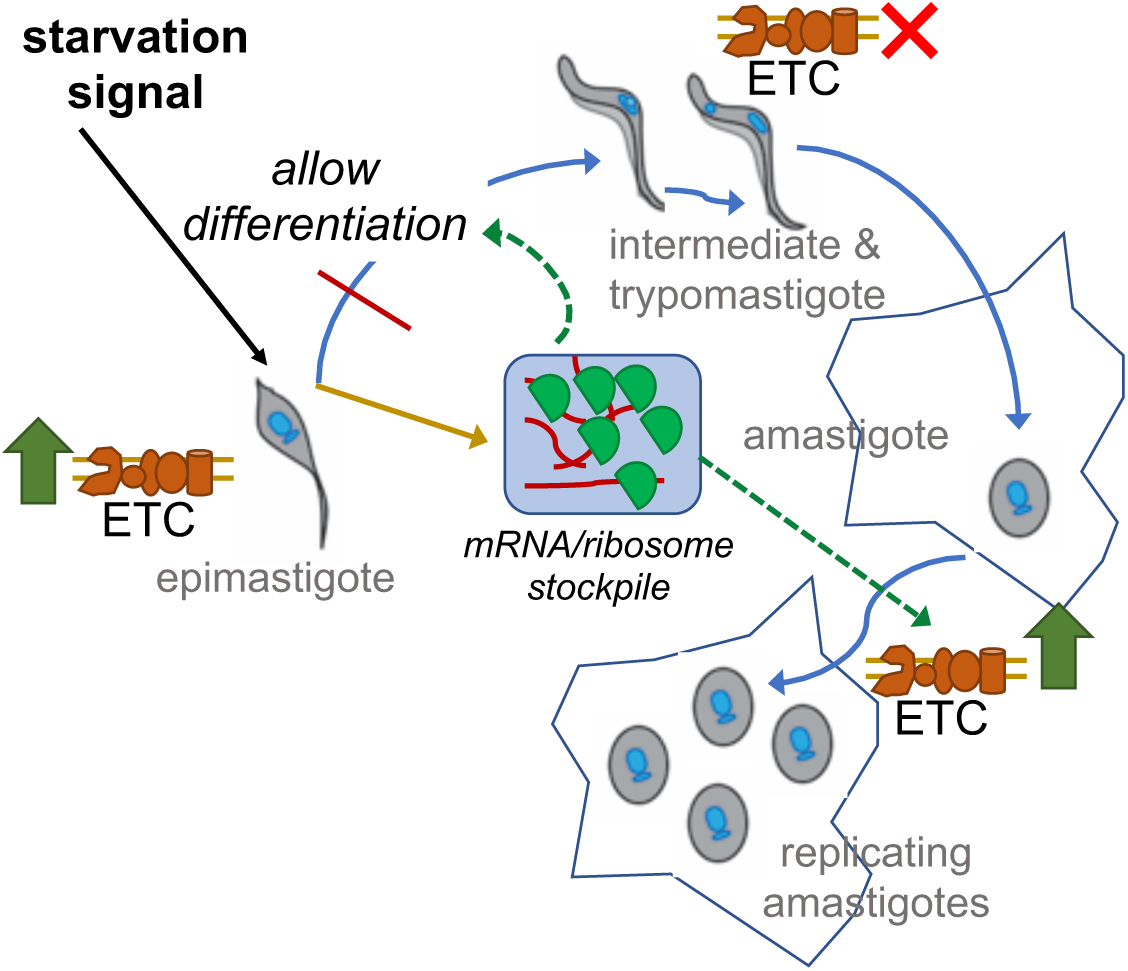
Some potential functions of starvation-induced spike in mitochondrial gene expression in *Trypanosoma cruzi*. The increase in mitochondrial RNAs upon starvation could serve as a retrograde signal to the parasite that it is competent for differentiation, as the cell has an adequate stockpile of material in the mitochondrion that will allow for restoration of the ETC and its activity once infectious cells have transformed into amastigotes and resumed replication. Grey text, parasite life stages; ETC, electron transport chain.

Alternatively, there are other explanations for our results. There could be minor changes in ETC function in *Tc*RND overexpressing cells resulting from depletion of edited mRNAs that do not affect normal epimastigote growth, but during differentiation are significant enough to impact starvation-compromised respiration. This is a valid alternative but hard to link to the change 13RND1 and T7RND parasite length. Additionally, as nuclear-encoded ETC subunits are not changed during starvation when select mature mRNAs increase in abundance (Shaw *et al*., 2016), the link between mitochondrial mRNA and ETC complex levels has not been established. We also cannot rule out that small amounts of overexpressed *Tc*RND are aberrantly localized, leading to nonspecific effects in the cytosol. However, in that case it would be surprising that phenotypes of *Tc*RND off-target enzyme activity would be so specific. Ultimately, it will take considerable experimental effort to parse mechanisms underlying the *Tc*RND overexpression phenotype. But clearly, mitochondrial retrograde signaling is a very underexplored topic of molecular parasitology.

## Experimental Procedures

### Parasite culture

*T. cruzi* CL Brener (DoCampo laboratory, University of Georgia) and Sylvio X10 (ATCC® 50823) were maintained in liver infusion tryptose (LIT) medium (Castellani *et al*., 1967) supplemented with 10% fetal bovine serum (FBS) at 27 °C, 5% CO_2_ and passaged every 2-4 days except for the ribosome profiling studies for which CL Brener was passaged every 8 days. For those experiments, 48 h prior to gradient fractionation, 7.5×10^8^ cells were washed in TAU medium (190 mM NaCl, 8 mM phosphate buffer pH 6.0, 17 mM KCl, 2 mM CaCI_2_, 2 mM MgCI_2_) (Contreras *et al*., 1985) and resuspended in 50 mL of TAU, or 2.5×10^8^ cells were diluted to 50 mL to grow exponentially until harvest.

For metabolic flux analysis, cultures in LIT medium were initiated at a concentration of 1×10^6^ cells ml^−1^ incubated for 8 days for stationary phase starvation (D8 LIT). Severe restriction (48H TAU) and regular growth medium (D0 LIT) cultures were initiated 48 h prior to experimentation. For 48H TAU, 2×10^8^ cells were collected, washed and resuspended in TAU in a final concentration of 1×10^7^ cells ml^−1^. D0 LIT cultures were initiated with 2×10^6^ cells ml^−1^. For growth curves, cultures were started at 2×10^6^ ml^−1^ and counted every other day for eight days. Tet was added fresh every 24 h. In all experiments, a concentration of 5 mg ml^−1^ tet was used for induction.

### Molecular cloning and transfection

The *Tc*RND nucleotide sequence was amplified by PCR from CL Brener strain genomic DNA using the oligonucleotides *Tc*RND-Fwd (5’-TTCAGTCGACT<u>GGATCC</u>ATGCTGCGTCGTGYTGGTGTC-3’) and *Tc*RND-Rev (5’-AAAGTCGGGTCTA<u>GATATC</u>TCAAGAAGAGGATCTTTCGTCAAAAAAG-3’) designed to amplify either allele (TcCLB.510743.90 or TcCLB.510659.219). The PCR product was cloned into pJET1.2 (Thermo Scientific) and excised at BamHI and EcoRV sites to be inserted in these sites in pENTR4 (Invitrogen), and then transferred to p*Tc*INDEX-GW (Alonso *et al*., 2014) by recombination using LR clonase II enzyme mix (Invitrogen) per manufacturer instructions to generate p*Tc*INDEX-GW-*Tc*RND.

CL Brener epimastigotes (3-5 × 10^7^ parasites ml^−1^) were first transfected with pLEW13 (Wirtz *et al*., 1999) or the derivative p*Tc*rRNA-T7Tet (Taylor and Kelly, 2006) using the Nucleofector 2b (Amaxa) X-014 setting: 10^8^ cells were washed with phosphate buffered saline and resuspended in 100 μl of transfection buffer (Pacheco-Lugo *et al*., 2017), mixed with ~30 μg of plasmid DNA and submitted to electroporation, after which cells were transferred into 15 ml of LIT medium with 15% FBS and diluted in a 24-well plate. After 24 h, geneticin (G418) was added at a concentration of 100 μg ml^−1^ to obtain semi-clonal cultures. The new “parental” cell lines containing episomal pLEW13 or p*Tc*rRNA-T7Tet were transfected with p*Tc*INDEX-GW-*Tc*RND (below), following the same protocol minus the dilution on plates, to generate *Tc*RND tet-inducible polyclonal cell lines 13RND1, 13RND2 and T7RND after 4 weeks of selection with 250 μg ml^−1^ of Hygromycin B.

### In vitro *differentiation*

Culture differentiation to the trypomastigote forms was induced as previously described (Shaw *et al*., 2016). Briefly, 1 ml of exponential-phase parasites in LIT medium was added into 10 ml of RPMI medium without FBS and left undisturbed for 8 days. As a supplemental method, a traditional approach involving successive incubations in TAU and TAU3AAG was also employed (Contreras *et al*., 1985). To quantify the differentiation rate, thin smears were generated from the supernatant (non-adherent cells, described in Shaw *et al*. (2016)), methanol-fixed, and stained with 3% Giemsa in Sorensen buffer (Riccha Chemical Company). Slides were viewed at 600X and images were collected. Cells were quantified manually using the cell counting plugin in the software ImageJ 1.52k (NIH, USA). Only parasites with a fully elongated nucleus and rounded kinetoplast at the posterior end were considered trypomastigote forms. One hundred cells were counted for each of 3 replicate cultures for determining differentiation efficiency for mammalian cell infections. At least 300 cells were counted per each of 3 biological replicates for *Tc*RND overexpression studies.

### *Mammalian infection with* T. cruzi

The AC16 (Davidson *et al*., 2005) cell line was maintained in Dulbecco’s Modified Eagle’s Medium (DMEM)/F-12 with 12.5% FBS and 1% penicillin-streptomycin, and the 3T3-L1 (ATCC CRL-1658) line was maintained in DMEM with 10% FBS and 1% penicillin-streptomycin. Mammalian cells were cultivated by weekly passage and incubated at 37°C in 5% CO_2_.

For infectivity assays, 3T3-L1 were plated in 6-well plates at 5×10^4^ cells per well, in DMEM + 10% FBS, while AC16 were plated at 2.5 × 10^4^ cells per well in DMEM/F-12 + 12.5% FBS and incubated for 24 h. Medium was removed and replaced with warm DMEM + 2% FBS medium. Cells were infected with *T. cruzi* parasites from the top fraction of 8 day RPMI culture (Shaw *et al*., 2016) at stated MOIs (MOI includes trypanosomes of all cell morphologies). Infected mammalian cells were incubated for 24 h after which extracellular *T. cruzi* were removed by rinsing 5X with warm DMEM + 2% FBS. Remaining mammalian cells were incubated for an additional 48 h. Mammalian cells were rinsed with DMEM + 2% FBS medium to remove any remaining *T. cruzi* before RNA isolation or staining.

At the time of RNA collection, 1-2 wells of infected cells were dried and fixed with 100% methanol, permeabilized with 1 ml IF-PERM per well for 15 minutes at room temperature, and stained with 3% Giemsa in Sorensen buffer. Stained cells were covered with water and photographed at 400X. *T. cruzi* intracellular amastigote quantification and percentage were determined by counting 100 mammalian cells in each of 3 replicates. Differences in number of cells infected and number of amastigotes per cell were analyzed using 2-way ANOVA, *α*= 0.05.

### RT-qPCR

RT-qPCR was performed and analyzed as described (Shaw *et al*., 2016), including use of the same two normalization genes, *PRF* and *TERT*. Total RNA was isolated from exponential-phase epimastigotes, mixed cultures resulting from the differentiation protocol, 48 h starvation treatment, infected or uninfected mammalian cells from a single largely-confluent well of a 6-well plate for experiments analyzing relative life stage abundances, or from exponential-phase epimastigotes for uninduced and induced *Tc*RND, pLEW13, and untransfected CL Brener (CLB-WT) cells for experiments analyzing *Tc*RND effects on editing. Primers utilized for detection of mitochondrial RNAs in strain CL Brener are those of (Shaw *et al*., 2016); those used to detect *Tc*RND and for work in Sylvio X10 are provided in Table 1. Unless stated otherwise, samples represent 3 biological replicates, collected on different days, and each measured in duplicate wells.

**Table 1.**
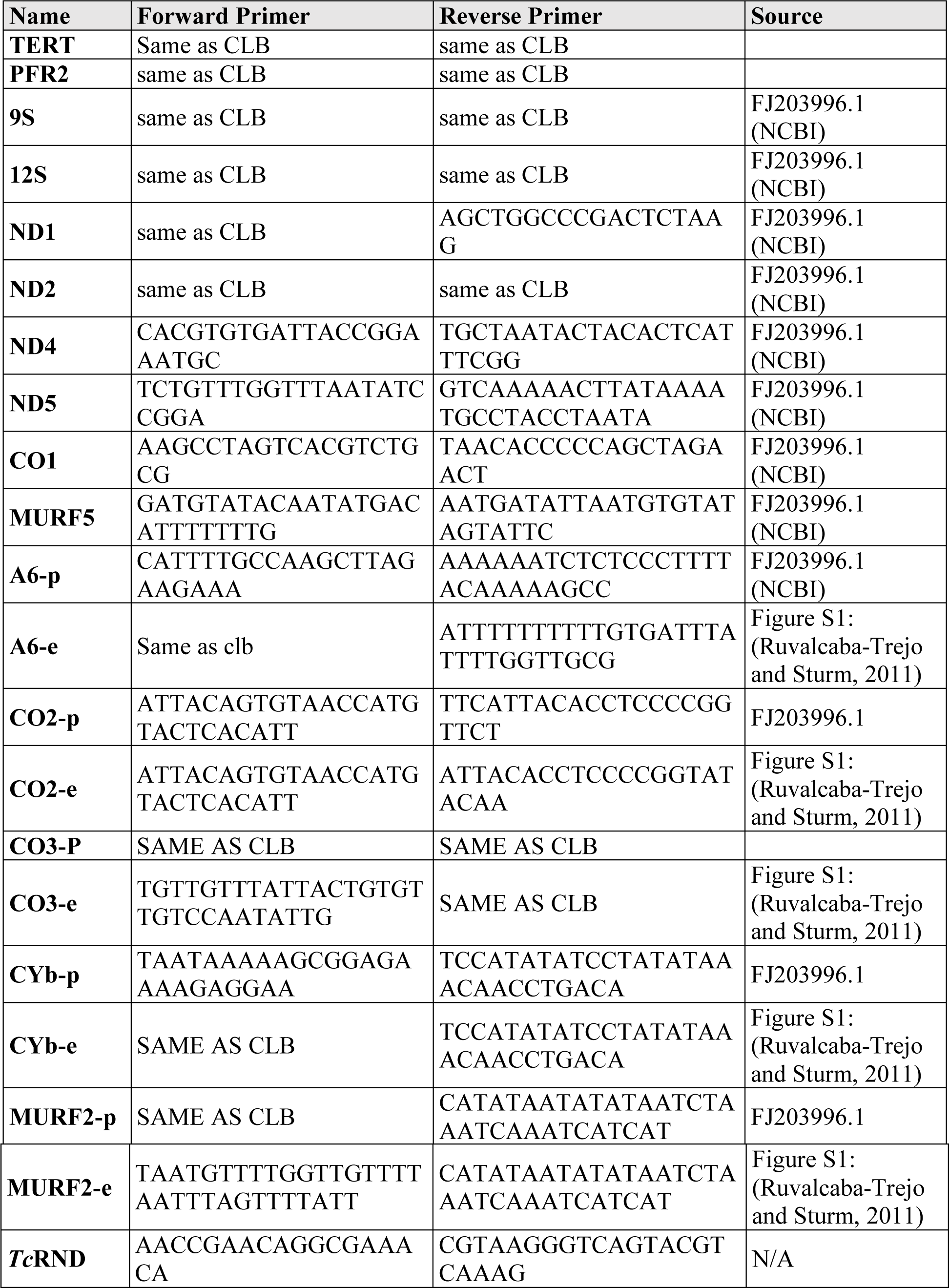
RT-qPCR primers specific to the *T. cruzi* Sylvio X10 strain, and those for detection of *Tc*RND in strain CL Brener. When the primer sequences between the strains are identical, the previously-described CL Brener primers were used (denoted “SAME AS CLB”). -p and -e indicates pre-edited and edited sequence, respectively. The last column indicates the source of the Sylvio X10 sequence used to determine specific primer sequences for each RNA.

### Ribosome and polysome profile analysis

Parasites were processed according to the protocol of Aphasizheva *et al*. (2016) with modifications. 5×10^8^ cells were washed in 5 ml of lysis buffer (125 mM KCl, 12 mM MgCl_2_, 50 mM HEPES, pH 7.6) and resuspended in 300 µl of lysis buffer with 1X cOmplete™ mini EDTA-free protease inhibitor (Roche). 10 µl of RNasin® (Ambion) was added, followed by 30 µl of 16% Triton X-100, and the cells were lysed with 30 passages through a 21-gauge needle. Four µl of TURBO™ DNase (Fisher) was added to the suspension and incubated on ice for 15 min. The extract was clarified by centrifugation at 21,000 × *g* for 15 minutes and 3 µl of 0.5 M EDTA was added. 30 µl of supernatant was reserved and diluted with 720 µl lysis buffer (input). The remainder of the supernatant was centrifuged as described and 15 750 µl gradient fractions were collected, from which RNA was collected by extraction with phenol:chloroform:isoamyl alcohol (25:24:1), pH 5.2. One µl of GenElute™-LPA (Sigma-Aldrich) prior to precipitation of the RNA with isopropanol at −20 °C overnight. Each fraction was resuspended in 10 µl H_2_O. For analysis of fraction rRNA content, 4 µl of every other fraction was electrophoresed on 6% acrylamide 8 M urea gels, except for the input lane, which contained approximately 1% of RNA from the initial lysis. After electrophoresis, RNA was transferred to BrightStar®-Plus (Ambion) membrane by semidry transfer in 0.5X TBE. After pre-hybridized in ULTRAhyb hybridization buffer (Thermo Fisher #AM8663), 5′ [γ-^32^P] ATP labeled oligonucleotide probes for rRNAs (GCTACATATATAACAACTGT (12S), CCGCAACGGCTGGCATCC, AAAAATTGGTGGGCAACA, and AATGCTATTAGATGGGTGTGGAA (all 3 identify 9S)) were added, and the membrane was hybridized overnight at 42 °C. After washing, blots were exposed to a phosphorimager screen and signal read with a BioRAD Personal Molecular Imager™. Quantification of ribosomal material per fraction was determined by densitometry, with pixel densities from 12S and 9S bands combined, and normalized to the same bands in the input lane for that experiment. The fractions were pooled into those containing nonassociated rRNA, small & large ribosomal subunits, monosomes, and polysomes by comparison with bacterial ribosome sedimentation coefficients.

Bacterial lysate was obtained using the small culture volume method from Mehta *et al*. (2012): 200 ml of *Escherichia coli* DH5α were grown to an optical density (OD_600_) of 0.6. The top 500 µl (71%) of the lysate was used for fractionation as with *T. cruzi* lysate. Fractions were then analyzed by electrophoresis on a 0.8% agarose gel and stained with ethidium bromide to observe bacterial rRNA.

### Respiratory capacity measurements

Prior to analysis, cells were collected and incubated in the dark, on ice, in 1 ml LIT or TAU with 15 µg/ml propidium iodide (Sigma). Cells were counted with brightfield, then counted again using the red fluorescence channel to identify inviable cells. For each condition, a minimum of 20 cells were counted and a minimum 95% viable cells was ensured.

Oxygen consumption rate (OCR) was measured with a SeaHorse XF^e^96 extracellular flux analyzer (Agilent) using the SeaHorse Mito Stress Kit (Aligent) as performed previously (Shah-Simpson *et al*., 2016; Kalem *et al*., 2018). Results were normalized staining the amount of cellular DNA in each well using SYBR Green I fluorescent dye (Invitrogen) as previously performed (Kalem *et al*., 2018) with Wave Desktop v 2.4 (Agilent) using Aligent’s recommended data analysis procedures. Six technical replicate wells were measured from each of four biological replicates. One-way ANOVA with multiple comparisons was performed to examine the significance of differences between strains and growth conditions.

### Cell length measurements

Thin smears were performed with exponential-phase epimastigotes cultures and stained with 3% Giemsa as described for *in vitro* differentiation. Total length and free flagellum length of one-hundred cells from each of the three biological replicates of each cell line were measured using NIS-Element software (Nikon Instruments, Inc).

### T. cruzi *crude subcellular fractionation*

Subcellular fractionation was performed using the hypotonic lysis procedure (Panigrahi *et al*., 2009) with modifications: 2.5×10^8^ cells were washed with cold NET (150 mM NaCl, 100 mM EDTA, 10 mM Tris-HCl, pH 8.0) and centrifuged at 1000 *g* for 10 min. The cells were then resuspended in 1.4 ml DTE (1mM EDTA, 1 mM Tris HCl, pH 8.0; Sylvio X10 cells were resuspended in 0.7 ml DTE, 0.7 ml DEPC treated water) and incubated on ice for 15 minutes. Cells were disrupted by 15 passages through a 25-gauge needle and immediately 190 µl of 60% sucrose was added. After taking a 100 µl aliquot (total protein fraction) lysate was pelleted at 15,000 *g* for 10 min at 4 °C. Supernatant was separated (non-mitochondria fraction) and the pellet was resuspended in 500 µl PBS (mitochondrial enriched fraction). 5 µl of total protein fraction, 5 µl of mitochondrial enriched fraction, and 10 µl of non-mitochondria fraction were used for subsequent immunoblotting.

### Tc*RND antibody purification and immunoblotting*

*Tc*RND-identifying antibodies were purified from serum from rabbits exposed to *Tb*RND (Zimmer *et al*., 2011). Antigen *Tc*RND was expressed in *E. coli* by cloning the open reading frame of *Tc*RND into pET42a (Novagen), and the mostly insoluble recombinant *Tc*RND expressed upon IPTG induction was separated from most soluble bacterial proteins by simple centrifugation, subject to SDS-PAGE, transferred to nitrocellulose, and used to affinity-purify *Tc*RND antisera from 2 ml of total serum. The purified *Tc*RND antibody preparation was used in a 1:500 dilution overnight at 4 °C, with goat anti-rabbit IgG as secondary antibody, at a 1:15,000 dilution (Li-Cor Biosciences). An Odyssey® Fc Imager (Li-Cor Biosciences) was used to detect the near-infrared signal from the secondary antibody. The *Tc*RND purified antibody was validated by its co-migration with a band on a ladder of approximately 38 kDa (Chameleon™ Duo Pre-stained Protein Ladder, Li-Cor Biosciences), by its cross-reactivity with native and over-expressed *Tb*RND in *T. brucei* (with the native *Tb*RND migrating at a similar position as the band detected in *T. cruzi*) and its specificity to a mitochondrial protein (Figure 6C and Figure S2).

*T. brucei* antibodies (Vondrusková *et al*., 2005; Fisk *et al*., 2010) were used to probe for MRP2 as a mitochondrial fraction control and PRMT1 as a non-mitochondrial fraction control; the *T. brucei* and *T. cruzi* MRP2 are ~65 identical and 75% similar in sequence and PRMT1 homologues are ~75 identical and 85% similar. Monoclonal anti-*α* tubulin antibody (Sigma-Aldrich T5168) was used as a loading control.

## Acknowledgements

We acknowledge the laboratory of Dr. Esteban Serra for the p*Tc*INDEX-GW plasmid and transfection advice, and Dr. Martin Taylor, London School of Hygiene & Tropical Medicine for the pT*cr*RNA-T7Tet plasmid. We thank Dr. Laurie Read for the *T. brucei* MRP2 & PRMT1 antibodies, RND antiserum, and the *T. brucei* RND wild-type overexpression cell line. This work was supported by American Heart Association Scientist Development Award 16SDG26420019 to SLZ.

## Author Contributions

RR and SLZ contributed to the conception and design of the study; RR, EKS, CKL, and SPF to the acquisition, analysis, or interpretation of the data; and RR and SLZ to the writing of the manuscript.

## Data Sharing

The data that support the findings of this study are available from the corresponding author upon reasonable request.

## Figures and Tables

**Figure S1.**
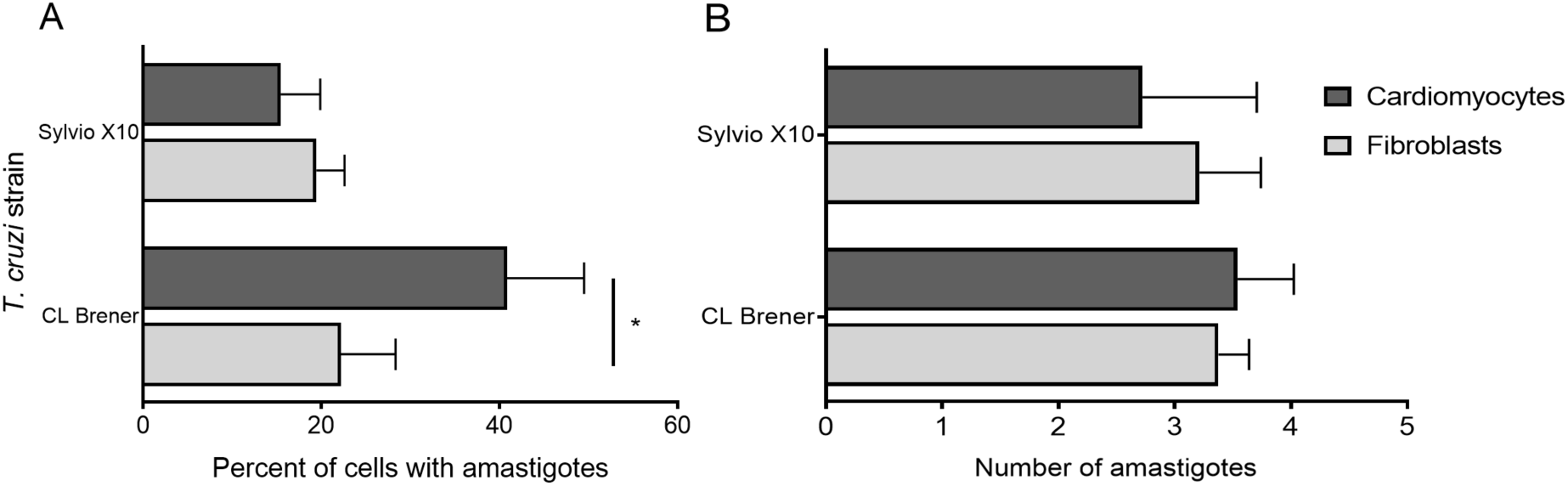
Quantification of *T. cruzi* infectivity assays in two mammalian cell lines. Infectivity assays utilized *T. cruzi* strains CL Brener or Sylvio X10 (y-axis) to infect cardiomyocytes and fibroblasts. **A**. Percentage of total mammalian cells that contain one or more *T. cruzi* amastigote within the cytoplasm. **B**. Average number of amastigotes in cytoplasm per infected mammalian cell. Data show mean of three biological replicates, with at least 100 mammalian cells counted for each replicate. Error bars represent standard error of the mean. * represents p *≤* 0.05

**Figure S2.**
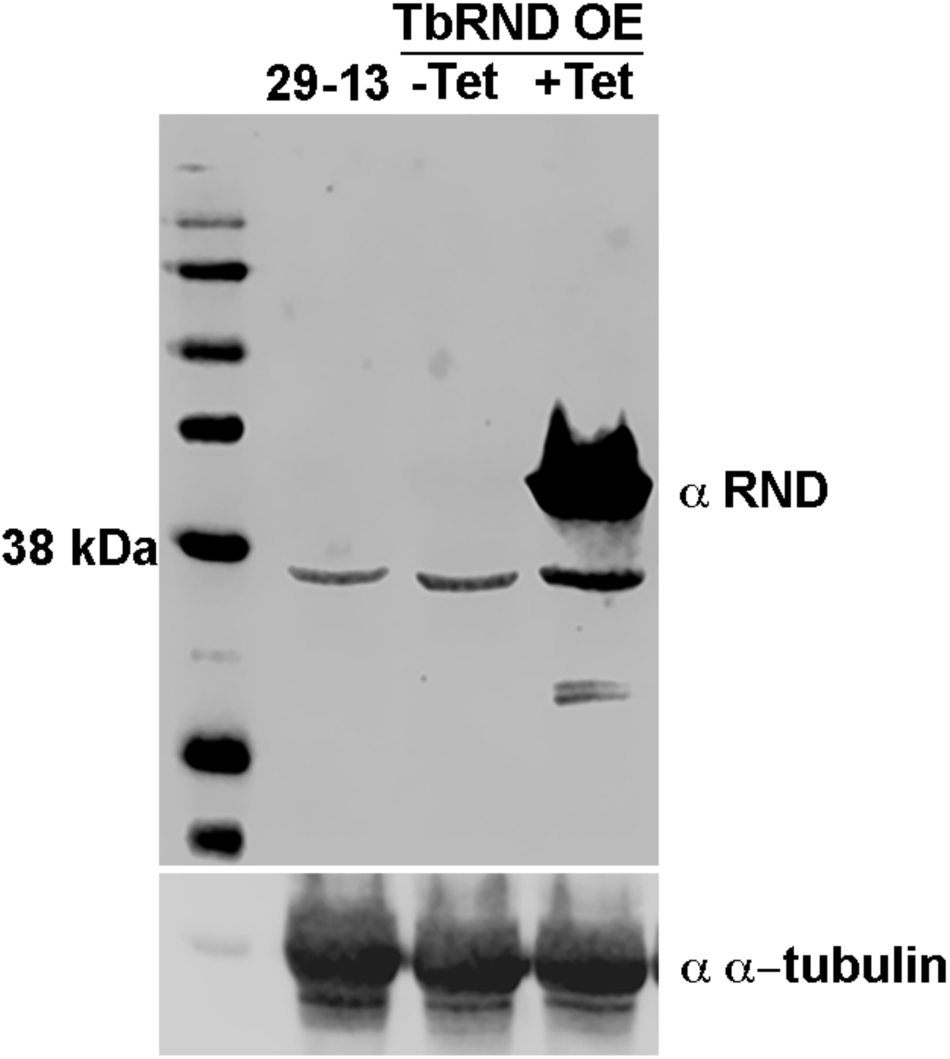
*Tb*RND can be detected with crude *Tb*RND antibody (serum) purified against recombinant *Tc*RND. *Tb*RND is identified as migrating as a single band near the 38 kDa ladder as expected for an approximately 42 kDa protein (including transit peptide). Loaded are the protein crude lysates from 1×10^7^ cells per lane from *Trypanosoma brucei brucei* cell line 29-13 (parent), and tet induced (48 h) and uninduced TbRND overexpressing cell line (Zimmer *et al*. 2011). TbRND that is overexpressed contains Myc, His, and Tandem Affinity Purification tags and appears as the upper band in the (+Tet) lane. Mouse anti-tubulin (*α*-tubulin, bottom panel) was used as a loading control.

**Figure S3.**
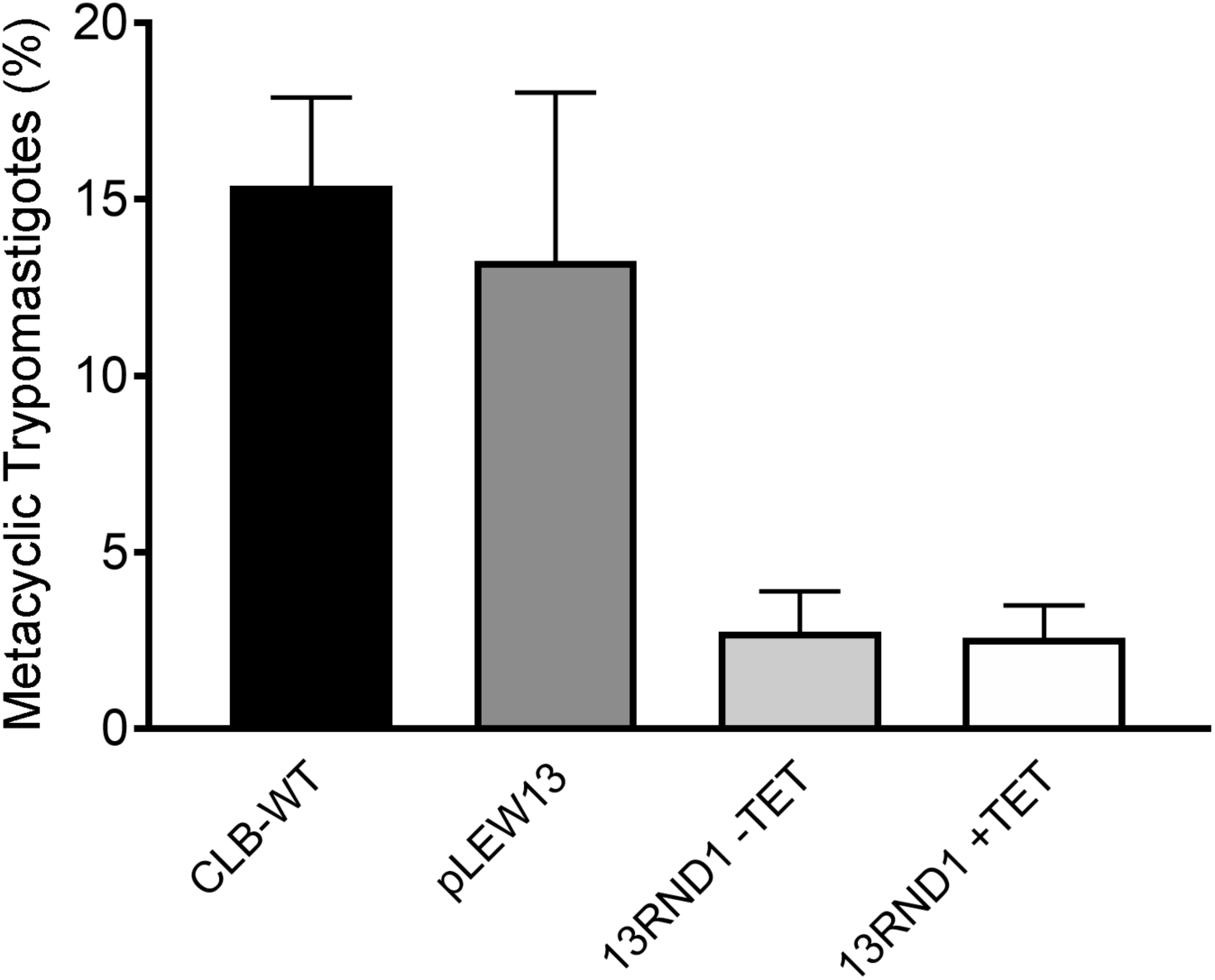
Phenotypic changes associated with *Tc*RND overexpression in *Trypanosoma cruzi*: Percentage of metacyclic trypomastigotes of wild-type strain CL Brener (CLB-WT), pLEW13-expressing parent strain (pLEW13), induced (+Tet) and uninduced (-Tet) *Tc*RND overexpressing parasites (13RND1) in culture differentiation utilizing TAU medium as detailed in Experimental Procedures. Error bars represent standard error of the mean. Experiment was performed one time by this differentiation method. Cultures were induced for 48 h prior to initiation of differentiation.

**Figure S4.**
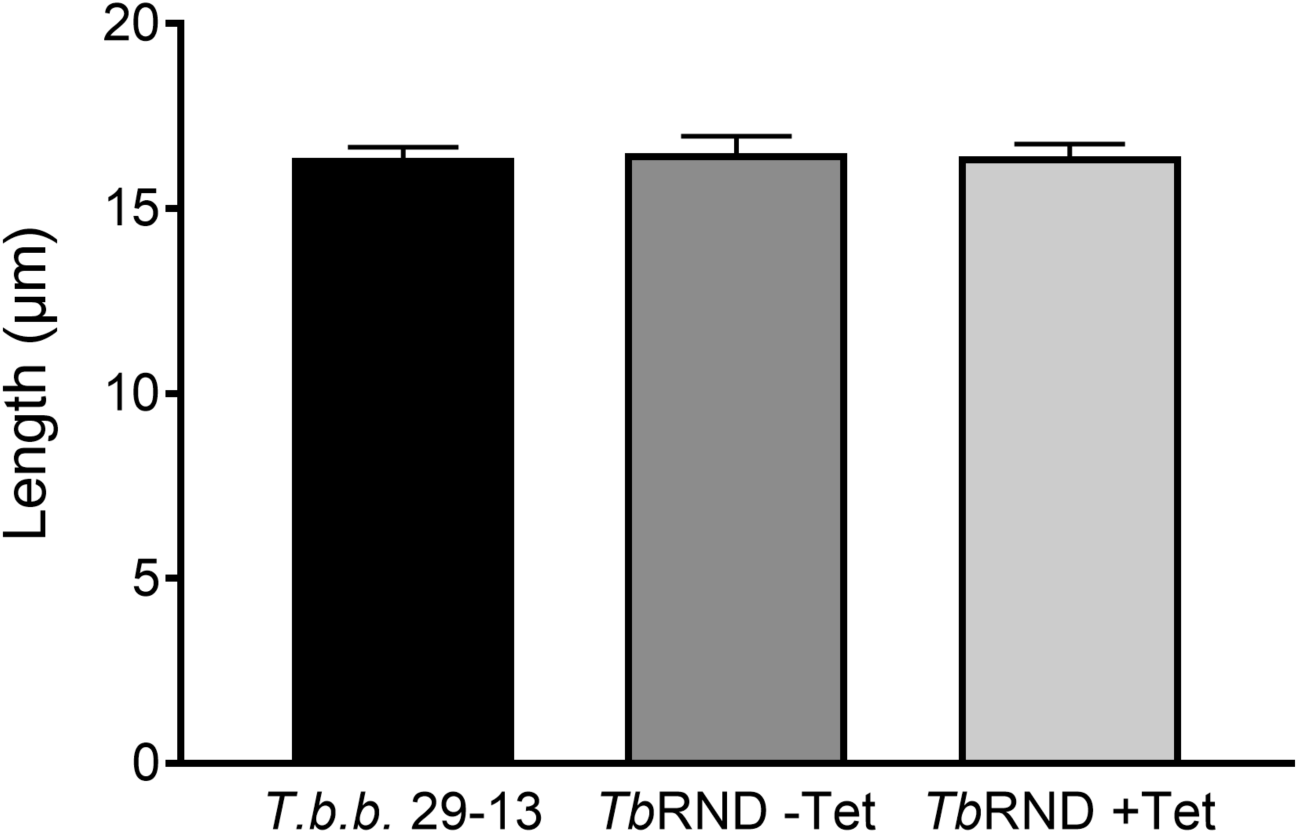
Total length of *Trypanosoma brucei* (*T.b.b*.) 29-13 strain, 48 h induced (+Tet) and uninduced (-Tet) *Tb*RND overexpressing parasites (Zimmer *et al*. 2011). At least 50 cells were measured. Error bars represent standard error of the mean. ANOVA P=0.9643.

